# Cell type-specific Interaction Analysis using Doublets in scRNA-seq (CIcADA)

**DOI:** 10.1101/2023.02.13.528326

**Authors:** Courtney Schiebout, Hannah E. Lust, Yina H. Huang, H. Robert Frost

## Abstract

**Motivation:** Doublets are usually considered an unwanted artifact of single-cell RNA-sequencing (scRNA-seq) and are only identified in datasets for the sake of removal. However, if cells have a juxtacrine attachment to one another in situ and maintain this association through an scRNA-seq processing pipeline that only partially dissociates the tissue, these doublets can provide meaningful biological information regarding the interactions and cell processes occurring in the analyzed tissue. This is especially true for cases such as the immune compartment of the tumor microenvironment, where the frequency and type of immune cell juxtacrine interactions can be a prognostic indicator.

**Results:** We developed Cell type-specific Interaction Analysis using Doublets in scRNA-seq (CIcADA) as a pipeline for identifying and analyzing biological doublets in scRNA-seq data. CIcADA identifies putative doublets using multi-label cell type scores and characterizes interaction dynamics through a comparison against synthetic doublets of the same cell type composition. In performing CIcADA on several scRNA-seq tumor datasets, we found that the identified doublets were consistently upregulating expression of immune response genes.

## 1. Introduction

Throughout the brief history of single-cell RNA-sequencing (scRNA-seq) analysis, instances where more than one cell has been recorded as a single-cell (known as a doublet for two cells or a multiplet for two or more cells) are assumed to be technical artifacts and are only identified during quality control processing in order to be removed from downstream analyses (Bais and Kostka, 2020; DePasquale *et al.*, 2019; McGinnis *et al.*, 2019). This is normally the correct course of action, as doublets can be an artifactual result of the cell dissociation protocol whereby two or more unrelated cells stick together (Bais and Kostka, 2020); however, the indiscriminate removal of all cells that appear to be doublets begs the question: is biologically relevant information lost when these doublets are discarded? In other words, are some doublets actually the result of physically interacting cells that do not successfully separate during tissue dissociation steps? If so, these doublets could provide novel information regarding juxtacrine cell-to-cell interactions within an experimental model, which is normally challenging to study using scRNA-seq data.

Interaction analysis is an ongoing area of research in scRNA-seq analysis, particularly in the case of juxtacrine interactions where the cells interact with one another by physical contact, given that the loss of cell location information due to dissociation makes it very difficult to determine which individual cells may have been involved in interactions. Numerous methods that have been put forth to address this challenge by adapting the results of interaction analysis to have lower resolution. Specifically, interaction analysis methods have been developed to give group- or cluster-level annotations, which identify interactions that are likely occurring between cells in whole clusters, such as SingleCellSignalR and CellChat (Cabello-Aguilar *et al.*, 2020; Jin *et al.*, 2021), or cell-level annotations, which identify when a certain cell has the expression profile befitting of sending or receiving a given interaction signal and can predict the other cell(s) involved but not identify them with complete certainty, such as NicheNet and ICELLNET (Browaeys *et al.*, 2020; Noёl *et al.*, 2021). These methods are relatively robust, allowing investigators to at least perform differential interaction analysis between experimental models. However, given that neither type of method allows for the full identification of both cells involved in an interaction, they are inherently limited in their specificity. Furthermore, these methods do not take into account any of the doublets or multiplets potentially present in the data, but rather remove or ignore them in their analysis. Especially in the case of juxtacrine interactions, maintaining and analyzing physically interacting cells in the form of multiplets could be extremely informative for cell-cell interaction analysis. Some researchers have recently explored the deliberate partial dissociation of tissues to enable the transcriptomic profiling of both singlets and interacting doublets (Boisset *et al.*, 2018; Halpern *et al.*, 2018). To date, these approaches have been limited to the identification of interacting cell types from the merged expression profile of the doublet. Although an improvement on earlier techniques, valuable information in the transcriptomic profile regarding the phenotypic impact of the interaction is ignored.

To address the challenges of interaction analysis using scRNA-seq data, an increasing number of interaction analysis methods have been developed that target spatial transcriptomics data, which maintains information on tissue architecture (Chen *et al.*, 2015; Ståhl *et al.*, 2016; Dries *et al.*, 2021; Cang and Nie, 2020). These allow cells in close proximity to be identified as such and analyzed with this context in mind. These methods are ideal for interaction analysis given their focus on maintaining cell location while performing transcriptomics, and serve as a useful gold standard for validating interactions inferred from methods like scRNA-seq that do not preserve location information; however, given that many datasets already exist in the scRNA-seq space that lack spatial information or validation, developing a method that can be used in these cases is a necessary and useful pursuit.

Here, we present Cell type-specific Interaction Analysis using Doublets in scRNA-seq (CIcADA), a method for discerning biologically relevant doublets and quantifying their interactions in scRNA-seq. We accomplish this by analyzing a joint scRNA-seq/Cellular Indexing of Transcriptomes and Epitopes by Sequencing (CITE-seq) dataset from the B16F10 model of mouse melanoma that was intentionally partially dissociated in order to preserve potentially bound cells. This allowed us to investigate doublet identification in a dataset we were confident contained doublets and whose experimental model was of interest for immune cell interactions. We then further tested our results with publicly available scRNA-seq data to illustrate the robustness of our findings on datasets where doublets were not intentionally maintained. We further validated our approach with Visium (Chen *et al.*, 2015) spatial data from the B16F10 model that recapitulates the findings from the scRNA-seq data. Throughout our investigation, we find that biologically relevant doublets can indeed be identified and that their resulting interactions are potentially clinically significant.

## 2. Methods

### a. CIcADA method

The details of the CIcADA method are described in-depth in the following methods sections, but in brief, CIcADA carries out the following steps (and visualized in Figure 1):

1. Perform cell type scoring on each of the cells in an scRNA-seq dataset using CAMML or ChIMP (depending on absence or presence of joint CITE-seq data) and identify doublets from cell types with notable overlap in the data
2. Create synthetic doublets from high-confidence singletons to serve as a comparison for the detected doublets
3. Cluster the synthetic and true doublets and contrast them using differential gene expression analysis

**Figure 1.**
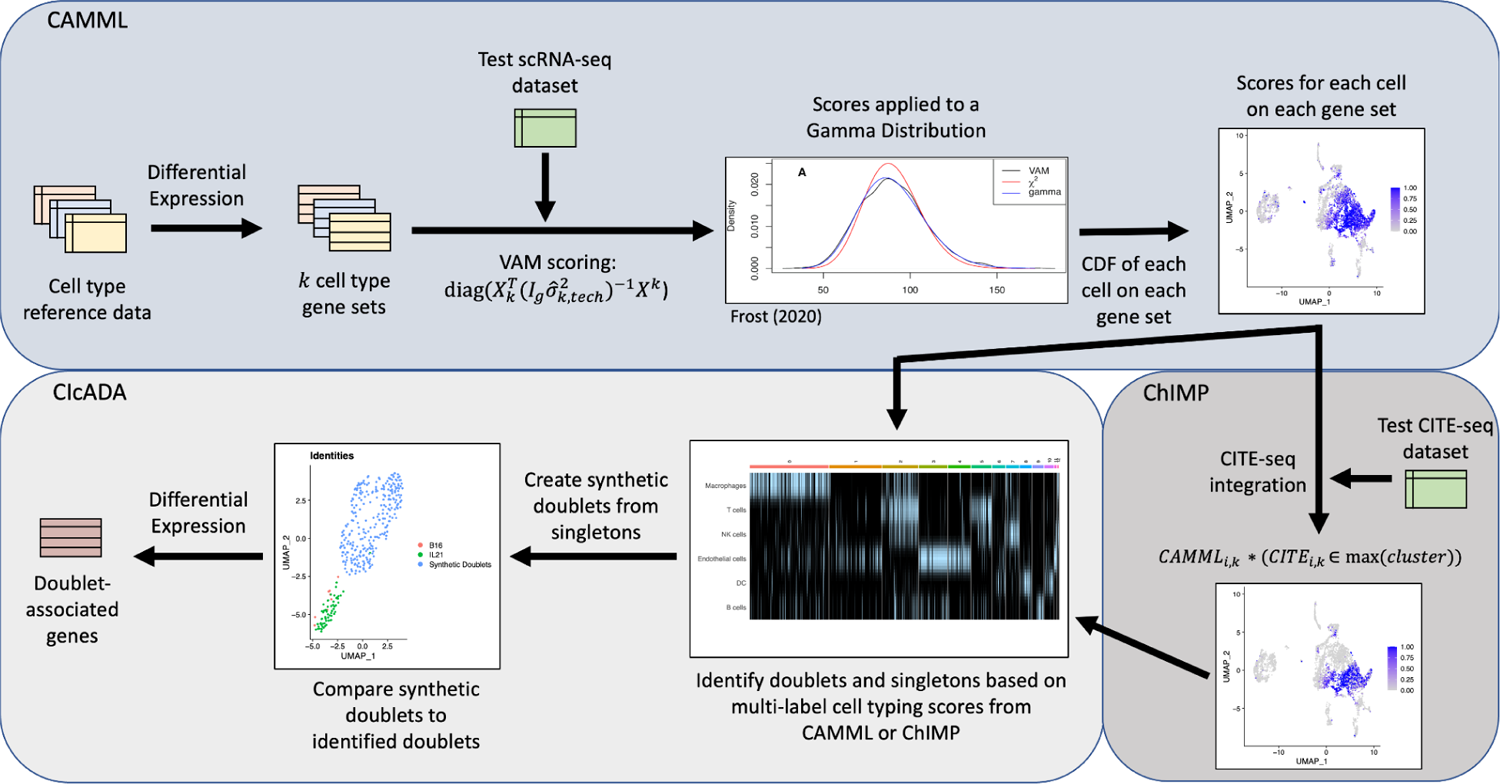
Flowchart of cell type and doublet identification steps.

This pipeline is explored using three datasets: a joint scRNA-seq/CITE-seq dataset from WT B16F10 and B16F10 IL-21 mouse melanoma with complementary Visium spatial transcriptomics data, a joint scRNA-seq/CITE-seq dataset from a MALT tumor available on 10X Genomics (10k Cells from a MALT Tumor, 2018), and a scRNA-seq dataset comprised of lymphoma tumor cells also available on 10X Genomics (Hodgkin’s Lymphoma, Dissociated Tumor: Whole Transcriptome Analysis).

#### i. Identify doublets with CAMML/ChIMP

To identify cells with a strong probability of being doublets, cell-type identification was performed using the ChIMP (CAMML with the Integration of Marker Proteins) method (Schiebout and H Robert Frost, 2022) on datasets containing joint scRNA-seq and CITE-seq data. For the datasets lacking CITE-seq data, cell-typing was performed using the CAMML (Cell-typing using variance Adjusted Mahalanobis distances with Multi-Labeling) method (Schiebout and H. Robert Frost, 2022). CAMML performs multi-label cell typing of scRNA-seq data by using the Variance-adjusted Mahalanobis (VAM) method to score each cell in the dataset for a collection of cell type-specific weighted gene sets (Frost, 2020; Schiebout and H. Robert Frost, 2022). The generated gene set scores, which are bound between 0 and 1, capture the association between each cell and the cell types of interest. ChIMP builds upon this process by adding an integration step for CITE-seq data whereby the CITE-seq counts for a given cell-type marker are separated by k=2 k-means clustering (Schiebout and H Robert Frost, 2022). Cells with CITE-seq counts in the higher value cluster keep their CAMML score for the complementary cell type, while cells with CITE-seq counts in the lower value cluster are scored as zeroes (Schiebout and H Robert Frost, 2022). This CITE-seq integration step makes ChIMP more confident but also more conservative than CAMML. A workflow of CAMML, ChIMP, and CIcADA is visualized in Figure 2.

**Figure 2.**
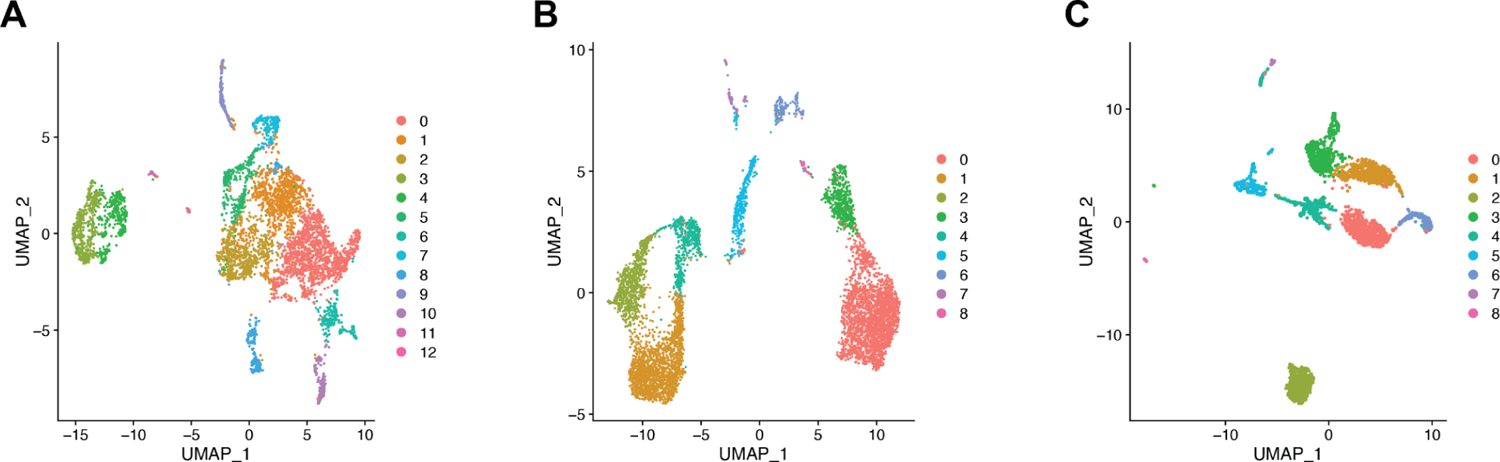
Visualization of the UMAPs and clusters for **A.** the B16F10 melanoma dataset, **B.** the MALT dataset, and **C.** the lymphoma dataset.

In each dataset for this investigation, the gene sets used for cell-typing were built with celldex, whereby differential expression analysis of cell types of interest is performed with edgeR with a one vs. all approach and genes with a log fold-change of greater than 5 are included in the relevant cell type gene set (Aran *et al.*, 2019; Robinson *et al.*, 2010). Cells are then scored with VAM, with genes weighted by their log2 fold-change value for that cell type. Cells that scored above a 0.75 for any two cell types were identified as potential doublets. Of note, this process inherently excludes the identification of homogeneous doublets in order to focus on interactions between differing cell types. Potential doublets were further filtered into groups based on the cell type composition, e.g., T cell/macrophage doublets, and the most notable doublet overlap was investigated. We did not attempt to identify and analyze higher-order multiplets given the small number of biological multiplets detected in the analyzed datasets, with fewer than 3% of cells in any dataset having more than 2 cell types with scores above 0.75.

#### ii. Separate artifactual and biological doublets

Once heterogeneous doublets have been identified, the next challenge is to determine if they are biological, i.e., cells undergoing a juxtacrine interaction, or just artifacts of the tissue and library preparation protocols used during scRNA-seq processing. Thus, a separate analysis step is needed to separate the heterogenous doublets into biological and artifactual groups. To realize this separation, the CIcADA method compares the expression profile of identified heterogenous doublets against synthetic doublets with the same cell type identities that are generated from singleton cells in the same dataset (the process for producing synthetic doublets is outlined in the following methods section). This procedure is carried out as follows: the raw counts from the identified heterogenous doublets and synthetic doublets are merged into a single scRNA-seq dataset, processed, and clustered to separate doublets according to their cluster identification. These datasets are processed using the same steps as outlined in the “scRNA-seq/CITE-seq data processing” methods section. For each unique cell type combination among the identified doublets, differential gene expression analysis is performed to identify transcriptomic differences between the biological and synthetic doublets.

#### iii. Run CAMML/ChIMP on singleton data and create synthetic doublets

To create synthetic doublets that represent a non-interacting benchmark for the separation of discern artifactual from biological doublets, we first identified high-confidence singletons in each scRNA-seq dataset using the cell type scores generated by ChIMP (Schiebout and H Robert Frost, 2022) or CAMML (Schiebout and H. Robert Frost, 2022). Specifically, if a cell scored above 0.75 for one of the doublet cell types of interest and no higher than 0.5 for any other cell type, it was defined as a confident singleton. These singleton cells were then combined with other singleton cells of the complementing cell type by summing their gene raw counts to produce synthetic doublets. For example, in the mouse B16F10 scRNA-seq/CITE-seq dataset, a population of macrophage and T cell doublets was identified by ChIMP (Schiebout and H Robert Frost, 2022). To create synthetic T cell/macrophage doublets, singleton T cells and macrophages were identified as cells in the B16F10 data that scored above 0.75 via the ChIMP method for either macrophage or T cell and no greater than 0.5 for any other cell type. These singletons were randomly combined by summing their raw gene counts to create synthetic T cell/macrophage doublets. Because both synthetic and artifactual doublets capture a combined transcriptome in the absence of juxtacrine interactions, we expect them to have similar mean expression profiles. While creating synthetic doublets in order to identify doublets in a dataset is not a new concept, prior methods do so without regard for the cell types present, inhibiting any investigation of their biological significance as it relates to specific cell-cell interactions (McGinnis *et al.*, 2019; Bais and Kostka, 2020).

#### iv. Separate experimental and synthetic doublets by clustering

The raw gene expression counts for the doublets identified in the experimental data and the synthetic doublets are built into Seurat datasets and merged (Satija *et al.*, 2015). Each object is filtered to retain cells with at least 100 genes and genes present in at least 50% of cells in that dataset. Following the merge of the datasets, mitochondrial and ribosomal genes are removed for clarity in downstream differential expression analysis. Following this, the count data is transformed by natural-log transformation and centered and scaled (Satija *et al.*, 2015). The data is finally clustered and visualized on 30 principal components with the same parameters as the original datasets, i.e., shared nearest neighbors clustering at a resolution of 0.25 and visualization of the first two UMAP dimensions (Satija *et al.*, 2015; McInnes *et al.*, 2018). Following this, the clusters are evaluated relative to original identities of the doublets, either found in the experimental data or synthetic, and only doublets in clusters entirely made of experimental doublets are considered to be biological doublets and kept for further analysis. This is an intentionally stringent approach for filtering in order to maintain the highest likelihood that the analyzed doublets represent interacting cells. The biological doublet clusters are then compared with synthetic doublet clusters using differential gene expression analysis to discern the meaningful biological impact of the potential juxtacrine interaction.

### b. B16F10 data processing

#### i. B16F10 and B16F10 IL-21 Tumor Induction and Tumor Infiltrating Leukocytes (TILs) Isolation

Six- to 8-week-old C57BL/6 were injected subcutaneously with 5×10^5^ B16F10 parental or B16F10 cells constitutively expressing IL-21 (B16F10 IL-21) on the right flank. Tumors were excised, smashed, and filtered through a 70 μm cell strainer to create a cell suspension prior to separation on a 40% −80% percoll (GE) gradient. Viable tumor infiltrating leukocyte cell layers were collected for scRNA-seq and CITE-seq analysis. Single-cell RNA-seq and CITE-seq processing and transcript quantification was carried out using Cell Ranger from 10x Genomics (Zheng *et al.*, 2017). The following antibodies were used for CITE-seq: CD4, CD8b, CD3, CD45, NK1.1, Thy1.2, CD25, CD11b, CD19, CD11c, CD204, and IL-23R (antibody panel details are available in Supplementary Table 1).

#### ii. Visium 10X Genomics spatial transcriptomics data processing

Two B16F10 and two B16F10-IL-21 tumors were excised and separated from skin prior to embedding in Optimal Cutting Temperature (OCT), snap freezing cryomolds in liquid nitrogen, and storage at −80C. Tumors were cryo-sectioned by the Dartmouth Health Pathology Core facility and transferred to a Visium Spatial slide. The Visium Spatial slides were then aligned, processed, and quantified with Space Ranger from 10x Genomics (10x Genomics Space Ranger 2.0).

#### iii. scRNA-seq/CITE-seq data processing

The joint scRNA-seq/CITE-seq data collected from the B16F10 mouse model of melanoma was first merged to include data from both the WT and IL-21+ tumors. This merged data was then filtered to only include genes present in at least 10 cells and to only include cells with at least 100 genes. Cells with more than 5% mitochondrial genes were also excluded to avoid capturing damaged or dying cells. The scRNA-seq data was natural-log transformed and scaled (Satija *et al.*, 2015). The CITE-seq data was normalized by centered log ratio transformation and centered and scaled in the same manner as the transcriptomic data (Satija *et al.*, 2015). The scRNA-seq was then clustered by shared nearest neighbors with “FindNeighbors” and “FindClusters” in Seurat with a resolution of 0.25 and visualized with Uniform Manifold Approximation Projection (UMAP) on 30 principal components (Satija *et al.*, 2015; McInnes *et al.*, 2018). This resulted in 5,223 cells in 13 clusters with data for 16,248 genes.

#### iv. Spatial transcriptomics data processing

Four slides of tumors were collected and analyzed using the 10x Visium spatial transcriptomics platform (Ståhl *et al.*, 2016) for the same B16F10 model used to collect the scRNA-seq/CITE-seq data. Two slides were tumors of the wild-type (WT) B16F10 melanoma model and two were tumors of the B16F10 melanoma model with constitutive expression of IL-21. The resulting spatial transcriptomics data was processed and filtered using Seurat (Satija *et al.*, 2015). Each slides’ expression data was scaled and transformed using SCTransform on 3,000 variable features (Hafemeister and Satija, 2019). The data was then clustered with “FindNeighbors” and “FindClusters” in Seurat using a resolution of 0.25 on 30 principal components and visualized via UMAP (Satija *et al.*, 2015; McInnes *et al.*, 2018). After processing, the first WT slide had 12,967 genes across 3,107 spots and the second WT slide had 12,181 genes across 2,797 spots. The IL-21 slides had 12,670 genes across 2,189 spots and 10,292 across 1,263 spots, respectively.

### c. Confirmation of B16F10 T cell/macrophage interactions using spatial transcriptomics data

The spots for each B16F10 Visium slide were analyzed with multi-label CAMML to identify co-localization of cell types (Schiebout and H. Robert Frost, 2022). The gene sets used to estimate the cell type profile for each spot were built based on differential gene expression analysis from the ImmGen dataset in celldex on 6 cell types: macrophages, T cells, NK cells, B cells, dendritic cells (DCs), and endothelial cells, the same cell types used in the scRNA-seq analysis for the B16F10 model (Aran *et al.*, 2019). Genes that had a log2 fold-change of greater than 3 for a given cell type were included in the gene set for that cell type and weighted by the log2 fold-change value. The log2 fold-change cut-off was lowered for the Visium data relative to scRNA-seq data due to increased sparsity. CAMML was then run on each of the Visium slides to determine which spots represented a mixture of different cell types (Schiebout and H. Robert Frost, 2022). The CAMML generated cell type scores were then used to determine the rate of co-localization of potential doublet pairs and confirm doublet patterns identified in the scRNA-seq/CITE-seq data. Co-localization was defined as spots where each of the cell types of interest accounted for at least 10% of the sum of all cell-type CAMML scores for that spot (T cells or macrophages in this case). To determine if this rate of co-localization was greater than what would be expected for a slide, random permutations were carried out whereby the CAMML scores for each cell type in each spot were randomized without replacement and co-localization was recalculated on the randomized data. In other words, every CAMML score for T cells, for example, was randomly assigned to a single spatial spot, then the same was done for every other cell type. The true rate of co-localization was then compared to the distribution of 1,000 co-localization rates when CAMML scores were permuted. Moran’s I was also computed for each slide using a binary of the co-localization rate versus spot location.

## 3. Additional dataset processing

The same steps for scRNA-seq and CITE-seq processing performed on the B16F10 data was carried out for two additional freely available public scRNA-seq datasets from a Mucosa-Associated Lymphatic Tissue (MALT) tumor (10k Cells from a MALT Tumor, 2018) and a human lymphoma sample (Hodgkin’s Lymphoma, Dissociated Tumor: Whole Transcriptome Analysis), with some minor exceptions. Both datasets were sensitive to filtering by mitochondrial genes, and thus, to maintain cell counts, the cut-off for exclusion was set to cells with more than 10% of genes being mitochondrial. Additionally, the human lymphoma dataset did not include CITE-seq data, so the steps specific to processing that omics modality were skipped. Following these processing steps, the MALT data had 6,438 cells over 9 clusters with data for 15,518 genes and the human lymphoma data had 2,717 cells across 9 clusters with gene expression counts for 16,445 genes.

## 4. Results and discussion

### a. Doublet identification

ChIMP (or CAMML for the dataset of lymph node cells that did not have CITE-seq data) was used to perform multi-label cell typing of the analyzed single cell transcriptomics data for common immune cell types (Schiebout and H. Robert Frost, 2022; Schiebout and H Robert Frost, 2022). In the B16F10 model of mouse melanoma, ChIMP analysis was performed for 6 cell types: macrophages (CD11b+), T cells (CD4+ or CD8+), NK cells (NK1.1+), endothelial cells (CD45-), DCs (CD11c+), and B cells (CD19+) (Schiebout and H Robert Frost, 2022). Following ChIMP analysis, cluster 2 was identified as having a notable overlap of cell identities, particularly macrophages and T cells (Figure 2A). T cells and macrophages were therefore used to evaluate potential juxtacrine interactions that may have resulted in doublet formation. It was found that 69 cells met the cut-offs for cell-type co-expression required for classification as T cell/macrophage doublets. Of note, 61 of these 69 doublets originated from the IL-21+ data.

In the MALT dataset, ChIMP was executed for 4 cell types: B cells (CD19+), NK cells (CD56+), T cells (CD4+ or CD8+), and macrophages (CD14+) (10k Cells from a MALT Tumor, 2018; Schiebout and H Robert Frost, 2022). Some cell types used in the B16F10 ChIMP analysis were excluded from the analysis for this dataset due to the differing markers used for CITE-seq. In this dataset there was also a visible overlap in cell type scores between T cells and macrophages, particularly in cluster 4 (Figure 2B). When these cells were analyzed for scoring overlap, 385 cells were identified as potential T cell/macrophage doublets.

Finally, in the lymphoma dataset, the same 4 standard cell types used in the MALT ChIMP analysis were used for CAMML scoring: B cells, NK cells, T cells, and macrophages (Hodgkin’s Lymphoma, Dissociated Tumor: Whole Transcriptome Analysis; Schiebout and H. Robert Frost, 2022). As mentioned previously, this dataset does not have a paired CITE-seq dataset, so only scRNA-seq was used for cell typing. In this case, the most notable overlap of cell types was in cluster 5, where both NK cells and macrophages were highly represented (Figure 2C), with 155 cells identified as potential NK cell/macrophage doublets.

The cell pairs identified as potentially doublets are consistent with prior research on the tumor microenvironment. In particular, macrophages can be instrumental in activating both T cells and NK cells by releasing proinflammatory cytokines and can also be acted upon, in the form of activation or inhibition, by both T cells and NK cells (Ansell and Vonderheide, 2013; Bellora *et al.*, 2010; Zhou *et al.*, 2021).

### b. Doublet separation

Following identification of doublets using cell-typing, synthetic doublets were created for each dataset for comparison. In the B16F10 dataset, 279 high-confidence singleton T cells and 685 high-confidence singleton macrophages were identified. The number of unique synthetic doublets that can be created is limited by the size of the smaller cell type population, in this case 279 T cells. The raw count data for the synthetic doublets was then merged with the count data for the identified biological doublets and the merged dataset was normalized, clustered and visualized via UMAP using the same processing steps and with the same parameters as outlined in the “scRNA-seq/CITE-seq data processing” methods section (Satija *et al.*, 2015; McInnes *et al.*, 2018). UMAP visualizations of both the clustering of these doublets and their true identities is shown in Figure 3A.

**Figure 3.**
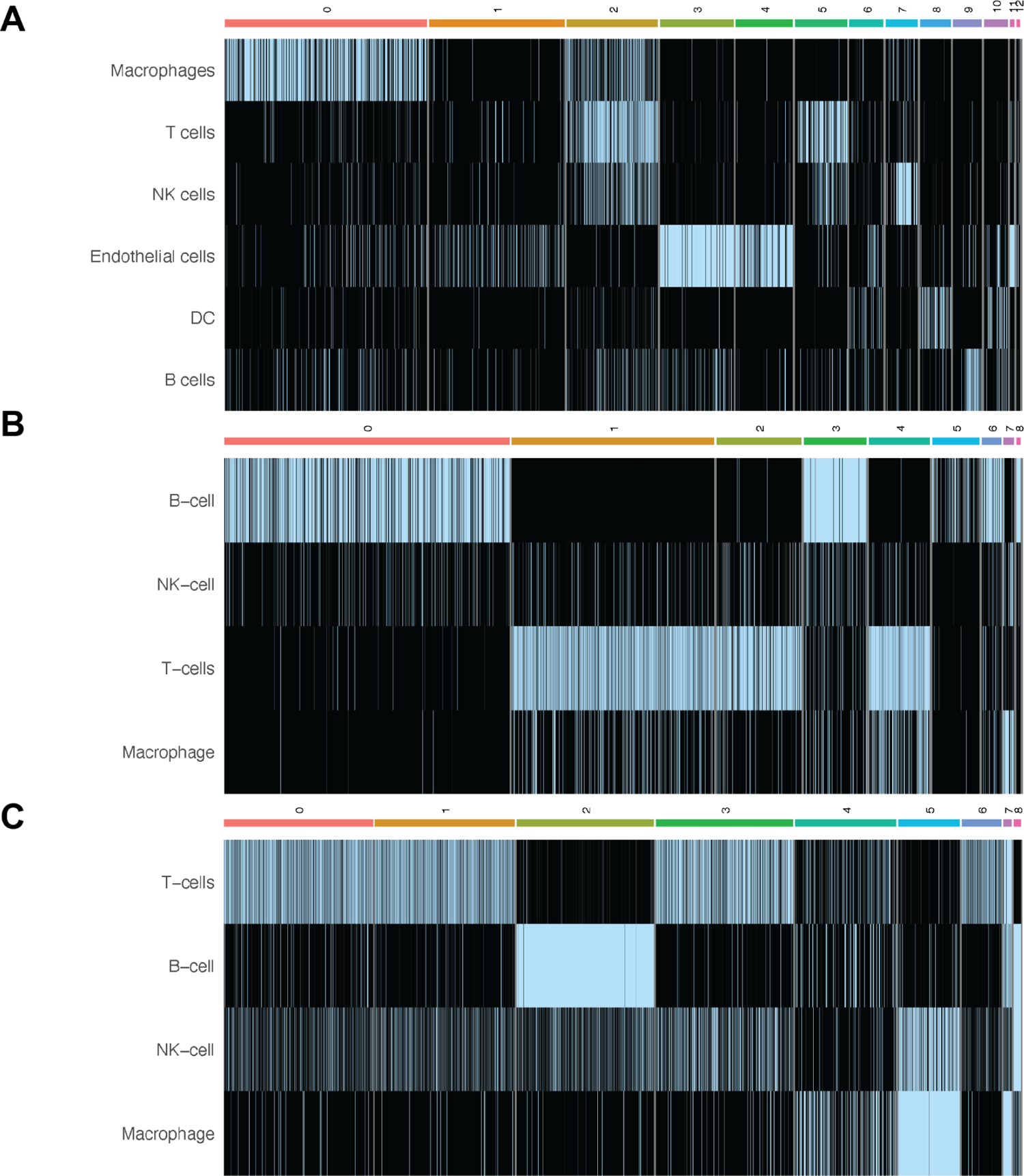
Heatmaps of the ChIMP/CAMML scores for each scRNA-seq dataset. **A.** The ChIMP cell type scores for the B16F10 data, with notable overlap of T cells and macrophages in cluster 2. **B.** The ChIMP cell type scores for the MALT data, with notable overlap of T cells and macrophages in cluster 4. **C.** The CAMML cell type scores for the lymphoma data, with NK cell and macrophage overlap in cluster 5. Columns correspond to the clusters identified in each dataset, which can be visualized in Figure 1.

The same synthetic doublet creation process was carried out for the MALT and lymphoma datasets (10k Cells from a MALT Tumor, 2018; Hodgkin’s Lymphoma, Dissociated Tumor: Whole Transcriptome Analysis). The MALT data had 1,459 high-confidence T cell singletons and 88 high-confidence macrophage singletons, resulting in 88 synthetic doublets. The clustering and original identities of these synthetic doublets and the doublets identified by cell-typing with ChIMP are visualized in Figure 3B. The lymphoma dataset had 39 high-confidence NK cell singletons and 158 high-confidence macrophage singletons, giving 39 synthetic doublets. These synthetic doublets clustered and visualized against the doublets identified with cell-typing by CAMML are visualized in Figure 3C.

As seen in Figure 3, most of the synthetic doublets and doublets identified by cell-typing with ChIMP or CAMML separate by clustering. This is not surprising given the stringent set of cut-offs used to identify both cell-typed doublets and high-confidence singletons and to include genes in the respective datasets. However, there are some cell-type specific doublets that do cluster with synthetic doublets, indicating that artificial doublets can be effectively identified should they pass through the aforementioned exclusionary steps. This highly stringent process for identifying doublets then allows us to evaluate how these doublet groups differ from one another and the biological implications of these distinct transcriptomic profiles.

### c. Doublet differential expression

Following the separation of doublets by clustering, resulting in the exclusion of any doublets that clustered with synthetic doublets, differential gene expression analysis was performed with “FindAllMarkers” in Seurat to compare the cluster of doublets identified by cell-typing with ChIMP to all other clusters present in the data (Satija *et al.,* 2015). In the B16F10 dataset, several genes with implications in immune response to cancer were upregulated in this group, including Cxcl9, interferon gamma, and Gpb2b (Pascual-García *et al.*, 2019; Jorgovanovic *et al.*, 2020; Arlauckas *et al.*, 2018, 1). A table of the most upregulated genes in each group is available in the Supplementary Information (Supplementary Table 2).

The differential expression analysis yielded similar results in the two validation datasets as well. The MALT dataset’s doublets had upregulation of immune response genes, including TXNIP and ETS1 (Chen *et al.*, 2020; Kim *et al.*, 2020, 1). The lymphoma dataset had genes related to NK cell and macrophage immunity, such as CXCL8 and MRC1, although interestingly, in this case, the roles of the upregulated genes represented both pro- and anti-tumoral immune responses (Montaldo *et al.*, 2012, 8; Walle *et al.*, 2022, 8; Arlt *et al.*, 2020, 206). Tables of all of the top upregulated genes in each of these datasets are available in the Supplementary Information (Supplementary Tables 2 and 3, respectively).

Collectively, these results indicate that utilizing multi-label cell-typing as a means of identifying potential biological doublets is effective and can provide important information regarding immune cell interactions with the tumor microenvironment. In all three cases, the genes found to be upregulated in the doublets identified using cell-typing, relative to synthetic doublets of the same cell types, are biologically relevant and informative for understanding how immune cells may be collaborating in response to cancer.

### d. Spatial transcriptomics validation

To further validate our findings, we utilized spatial transcriptomics (ST) data generated using the 10x Visium technology on the same B16F10 mouse model of melanoma analyzed via joint scRNA-seq/CITE-seq (Ståhl *et al.*, 2016). Specifically, the ST data was used to determine if the findings in the joint scRNA-seq/CITE-seq data could be detected in a different modality. We found a notable increase in the co-localization (as outlined in the “Confirmation with spatial transcriptomic data” methods section) of T cells and macrophages in the ST data generated on IL-21+ tumors, which supports the scRNA-seq findings where only 8 and the 69 biological doublets originated from the WT data (Figure 4). We further tested this finding with Moran’s I on the co-localization status of each spot. Every slide was significant for spatial autocorrelation with Moran’s I but the observed value for the IL21+ tumors was about an order of magnitude higher than the observed value in the WT tumors (Figure 4). Furthermore, following permutation of the CAMML scores, which serves to provide a permutation null distribution of the expected co-localization rates (see the “Confirmation with spatial transcriptomic data” section of the methods for details), we found that the rate of co-localization in the IL-21+ ST data was significantly higher than what would be expected by chance (according to a p < .05 on the empirical CDF of the permuted distributions). Interestingly, the WT tumors presented the opposite trend, where the rate of co-localization was actually lower than the average of the permuted distribution (Figure 5). Overall, these findings support the doublet analysis, with notable co-localization of T cells and macrophages, particularly in the cells originating from IL-21+ data.

**Figure 4.**
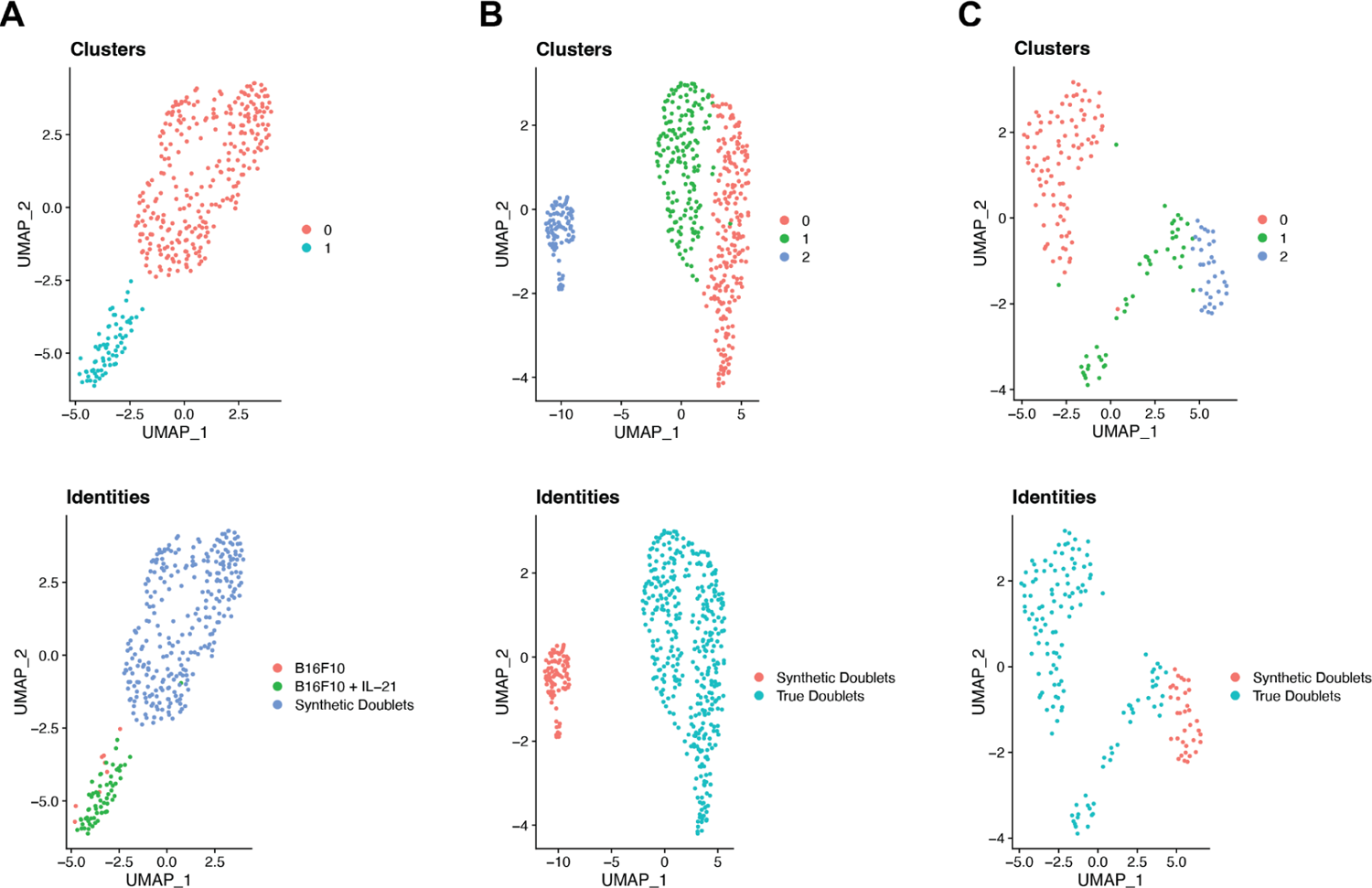
UMAP projections for each of the analyzed datasets (**A.** B16F10, **B.** MALT, and **C.** lymphoma) showing the clustering of true and synthetic doublets in the top row and their actual identities in the bottom row.

**Figure 5.**
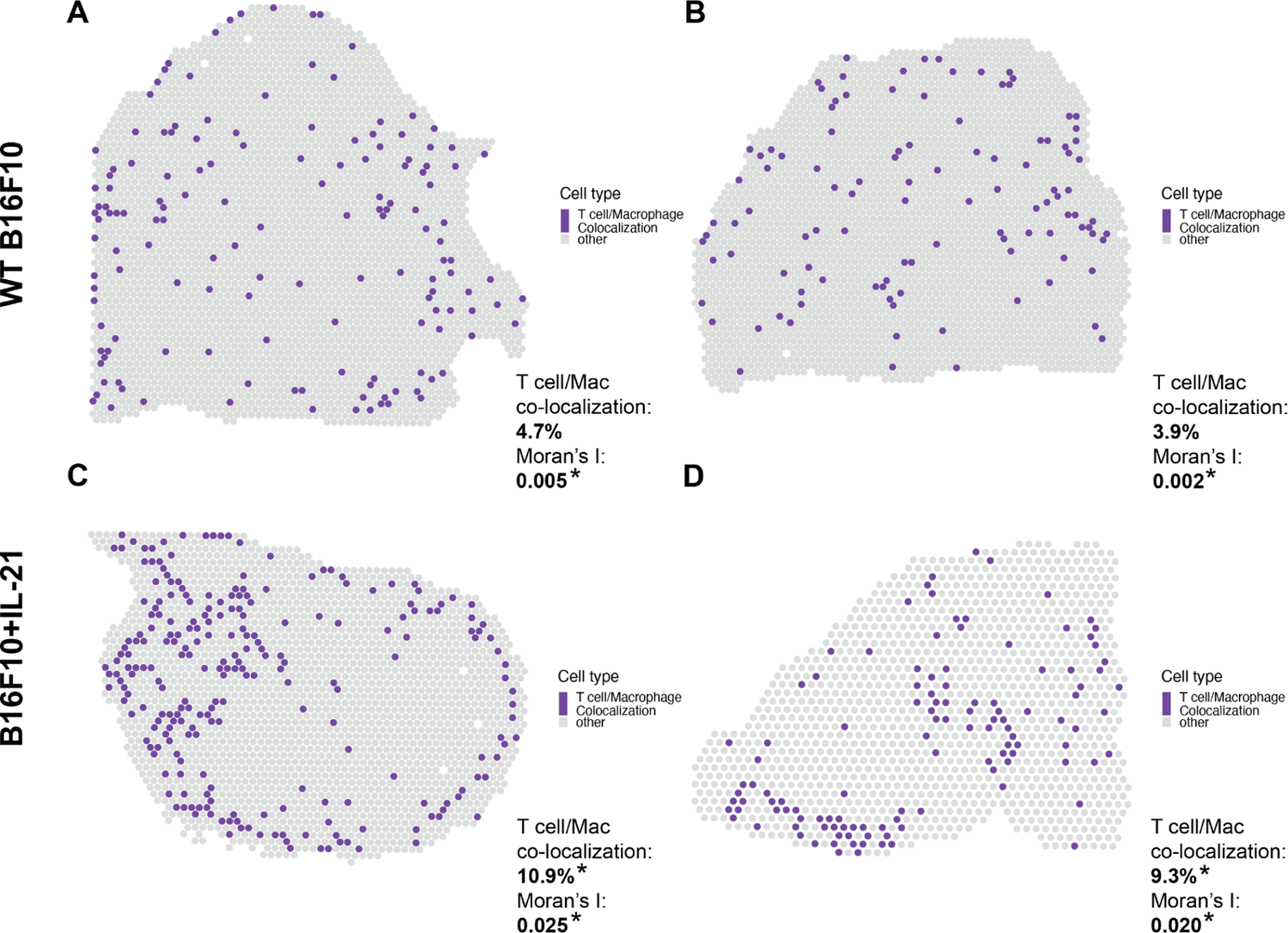
Spatial plots showing the proportions of T cell and macrophage CAMML scores at each Visium spot in the B16F10 data. Co-localization is the percent of spots where T cell and macrophage cell type scores each make up at least 10% of the score for that spot. Significance is determined based on the empirical CDF of random permutation of cell type scores across spots for that slide. Moran’s I observed values are also reported for each slide based on the spatial autocorrelation of the co-localization metric. The top two panels (**A** and **B**) are wild type B16F10 tumors and the bottom two panels (**C** and **D**) are tumors from B16F10 with constitutively active IL-21. The percent of spots where both T cells and macrophages scores make up at least 10% of all CAMML scores for that spot are provided in the bottom right of each panel.

To further validate the scRNA-seq findings, the upregulated genes found using differential expression analysis were evaluated using the ST data. Although some of the top genes could not be evaluated given the sparsity of ST data, Cxcl9 and Gbp2b both had nonzero ST expression data. In both cases, the average expression was significantly increased in spots that had co-localization of T cells and macrophages relative to spots that did not (Figure 6). Furthermore, the average expression of both genes was significantly increased in the IL-21+ tumor slides relative to the WT slides (Figure 6). These results further support the associations deduced from the scRNA-seq doublet analysis. Importantly, this validation is based on an independent transcriptomics modality, 10x Visium ST, that is particularly useful for evaluating cell-cell interactions since it maintains tissue architecture. Additionally, these findings recapitulate existing knowledge about the B16F10+IL-21 model of mouse melanoma, specifically that it has higher immune activation and smaller tumor size relative to the wild type model.

**Figure 6.**
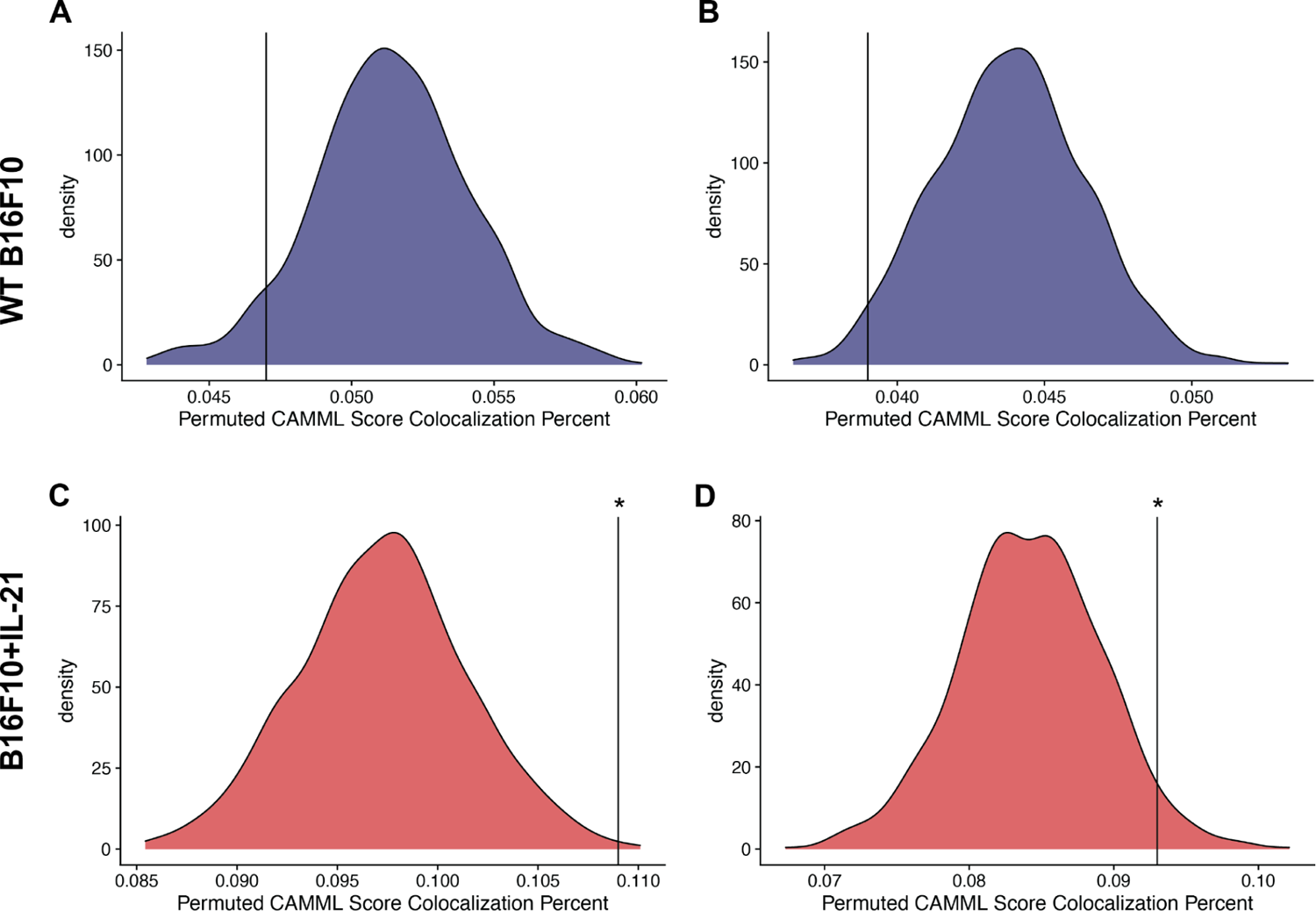
Density plots for the permuted distributions of co-localization statistics for each of the spatial transcriptomics slides: **A-B.** WT B16F10 slides 1 and 2 respectively and **C-D.** B16F10 + IL-21 slides 3 and 4. In each case, the black vertical line represents the true co-localization percent for that slide, with significance determined based on the empirical CDF of the permuted values.

**Figure 7.**
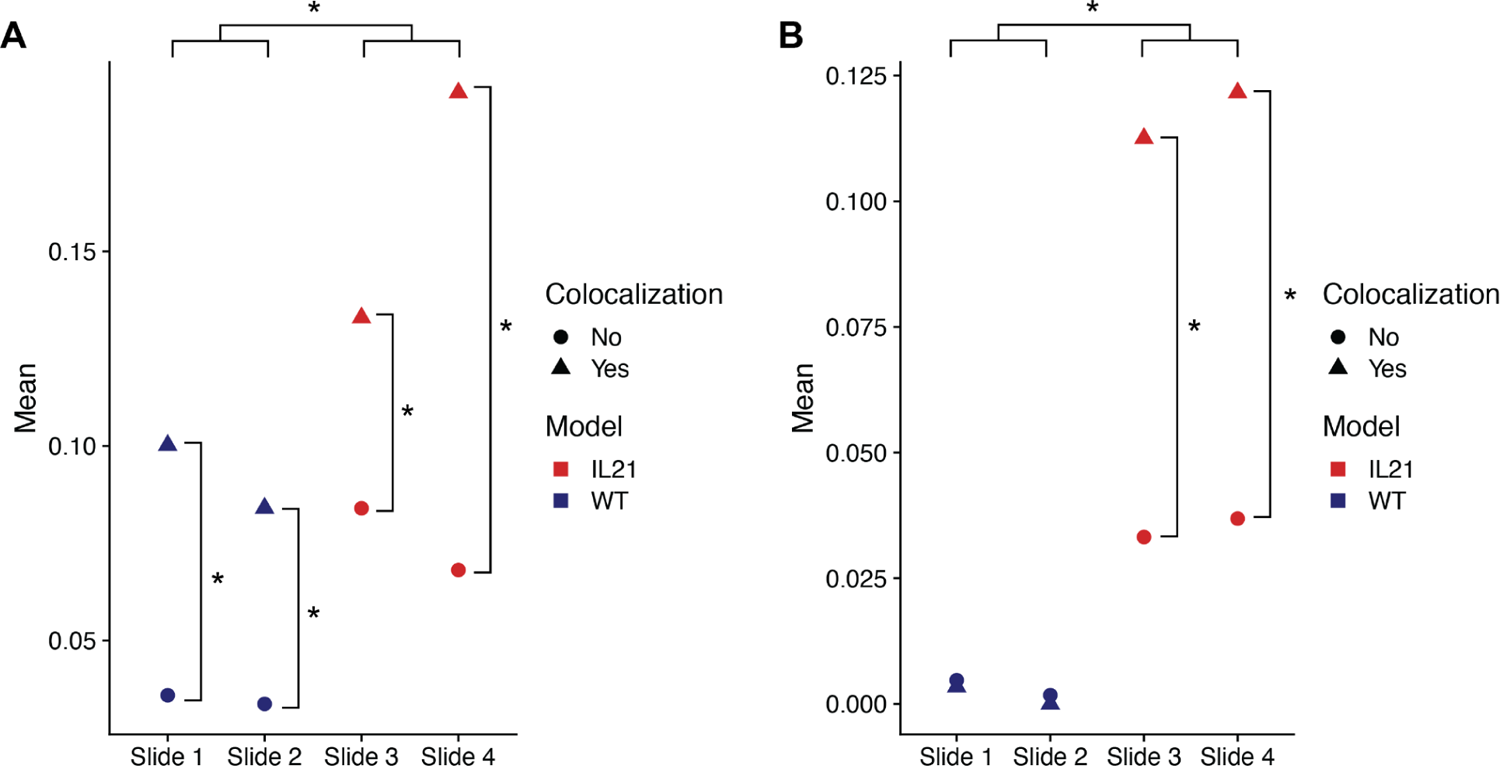
Average expression of the Cxcl9 and Gpb2b genes per spot in Visium slides from the B16F10 model of mouse melanoma. Points are split by colocalization status (true when macrophage and T cell scores account for at least 10% of all CAMML scores for that spot) and slide number. Slides 1 and 2 represent wild type B16F10 tumors and slides 3 and 4 represent B16F10 tumors with constitutively active IL-21, as designated by color. **A.** Average gene expression per spot of Cxcl9. **B.** Average gene expression per spot of Gbp2b. Significance between points was determined using the Wilcoxon Rank Sum Test.

## 5. Conclusion

Doublets occuring in scRNA-seq data have historically been treated as artifactual errors to be identified and removed before downstream analyses (Bais and Kostka, 2020; DePasquale *et al.*, 2019; McGinnis *et al.*, 2019). However, this approach overlooks the reason for doublet persistence in typical scRNA-seq dissociation protocols and their potential biological relevance. To evaluate scRNA-seq doublets, and to discover if they hold meaningful information regarding juxtacrine interactions, we developed the CIcADA (Cell type-specific Interaction Analysis using Doublets in scRNA-seq) method. CIcADA is a bioinformatics pipeline for identifying heterogeneous doublets in scRNA-seq data, separating doublets into artifactual and biological populations, and investigating the biological doublets for information regarding juxtacrine signaling within the source tissue.

We evaluated this method on a locally generated B16F10 scRNA-seq/CITE-seq dataset where only partial dissociation was carried out to improve the likelihood of maintaining interacting cell pairs. In this dataset, CIcADA successfully identifies single cell data points that have high doublet potential and, when compared to synthetic doublets, these candidate doublets showed high expression of genes associated with immune cell interaction in the tumor microenvironment. The findings from the scRNA-seq/CITE-seq analysis were validated using 10x Visium spatial transcriptomics data measured on the same murine B16F10 tumor model and the co-localization of cell types and gene expression analyses confirmed the signals detected in the scRNA-seq data (Ståhl *et al.*, 2016). Additional analysis of public scRNA-seq datasets that underwent the typical tissue dissociation process illustrates that our method is still effective at finding doublets with biologically relevant implications in cases where doublet maintenance was not a priority. CIcADA is a stringent method for detecting and evaluating biologically relevant doublets in scRNA-seq data, with a particular emphasis for utilization on scRNA-seq data of the tumor microenvironment.

## 6. Acknowledgments

This work was funded by National Institutes of Health grants R35GM146586, R21CA253408, P20GM130454 and P30CA023108. This work was also funded by the Burroughs-Wellcome Fund Big Data in the Life Sciences Training Program. We would like to acknowledge the supportive environment at the Geisel School of Medicine at Dartmouth College where this research was performed.

## Supplementary Information

**Supplementary Table 1.**
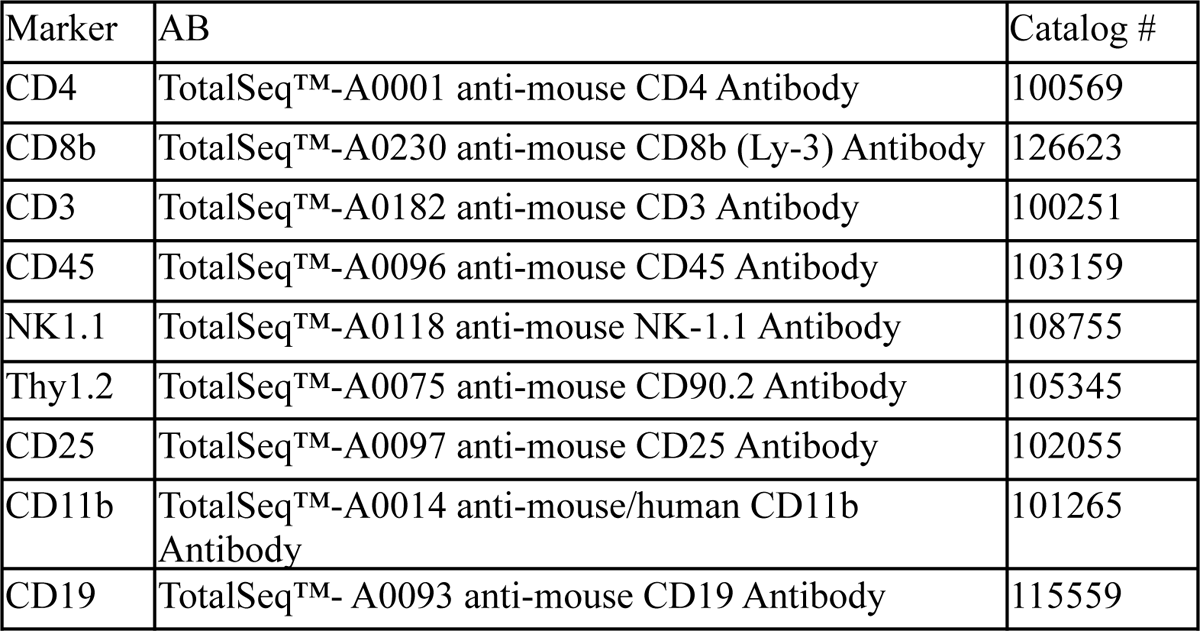
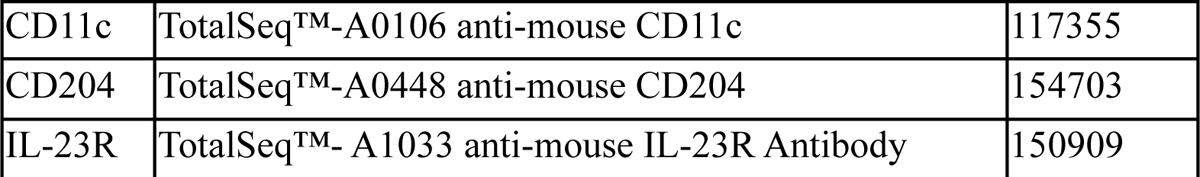
B16F10 mouse model of melanoma CITE-seq antibody panel.

**Supplementary Table 2.** Top 10 upregulated genes in the identified doublet clusters compared to the synthetic doublet clusters for each dataset.

| Dataset | Upregulated doublet genes |
| --- | --- |
| B16F10 (WT/IL-21) | Hmox1, Ccl5, Fcgrt, Klre1, Serpinb6b, Gpb2b, Cxcl9, Il18rap, Nr1h3, Ifng |
| MALT | ARID5B, RNF19A, TXNIP, TRAC, CD3G, TRBC2, ITM2A, CD2, ETS1, CD3D |
| Lymphoma | CXCL8, MRC1, LIPA, IL1RN, CLEC10A, EREG, C5AR1, G0S2, IL1B, F13A1 |

